# Birds have swing: Multifractal analysis reveals expressive rhythm in birdsong

**DOI:** 10.1101/157594

**Authors:** Tina Roeske, Damian Kelty-Stephen, Sebastian Wallot

## Abstract

Music is thought to engage its listeners by driving feelings of surprise, tension, and relief through a dynamic mixture of predictable and unpredictable patterns, a property summarized here as “expressiveness”. Birdsong shares with music the goal to attract and hold its listeners’ attention and might make use of similar strategies to achieve this goal. We here tested a songbird’s *rhythm*, as represented by the amplitude envelope (containing information on note timing, duration, and intensity), for expressiveness. We used multifractal analysis which is designed to detect in a signal dynamic fluctuations between predictable and unpredictable states on multiple timescales (e.g. notes, subphrases, songs). Results show that the songs' amplitude envelope is strongly multifractal, indicating rhythm is patterned by fluctuations between predictable and unpredictable patterns. Moreover, comparing original songs with re-synthesized songs that lack all subtle deviations from the “standard” note envelopes, we find that deviations in note intensity and duration significantly contribute to the rhythm’s multifractality. This suggests that birds render their songs more expressive by subtly modifying note timing patterns, similar to musical operations like *accelerando* or *ritardando*. Our findings bear consequences for neuronal models of vocal sequence generation in birds, as they require non-local rules to generate rhythm.

## 1. Introduction

### Structural analyses of birdsong

Birdsongs are complex vocal sequences unfolding with rich rhythmic and melodic structure across time. What determines the dynamic structure of a species’ song? Many structural analyses of song have focused on how acoustic characteristics of song represent *adaptations* to ecological conditions, for instance how a habitat’s specific sound propagation/attenuation properties affect song structure ^1–11^, or how structural aspects are affected by simultaneous singing of other hetero-or conspecific singers ^6^,^12–15^ or the energetic cost of producing different sounds ^16–19^. The structural aspects investigated were either high-level units like repertoires, song types, overall song/inter-song pause duration, or the occurrence of specific element categories such as trills, whistles, long, high-pitched, modulated notes ^6^,^20^, or of discrete types of notes ^13^.

Another line of research on birdsong structure has been concerned with the question what kind of algorithms the avian brain might use when generating note sequences, conceptualizing birdsong *syntactically*, i.e. as concatenations of distinguishable notes (“syllables”) whose transitions follow regular principles ^21^. Analysis inspired by formal language theory has modeled different species’ syntax in terms of simple first-order sequence generation in which current states only depend on just-previous states ^21^,^22^, or using slightly more complex sequence generation models that require a slightly longer memory ^23^.

Beyond the questions of ecological adaptation on the one hand and syntax generators on the other hand, we can take yet another perspective and focus on the proximate function of song to engage and attract a listener ^24^. Which aspects of song structure could be specifically designed to attract and hold the attention of conspecific birds? As this is a function that has been suggested to be shared by birdsong and music ^24^, research on human music might contribute relevant insight about what kinds of structures might be effective in fulfilling this goal.

Music psychology has put forth the hypothesis that what makes music attractive for listeners is its *dynamically fluctuating predictability* ^25^,^26^: That is, by building and breaking expectations on multiple timescales, music is thought to create a dynamic succession of different feelings. Stereotypic/predictable patterns result in a sense of fulfillment, relief, and ease of processing; unexpected/variable patterns in surprise, and delay of an expected pattern in tension ^26–28^. The constant fluctuation between variable/unpredictable and stereotyped/predictable patterns is believed to effectively attract and hold a listener’s attention.

We suggest using the term *expressiveness* for such fluctuations that may be used to engage a receiver’s attention^29^,^30^. In the present study, we ask whether there is evidence that birdsong uses similar mechanisms to attract its listeners, i.e. whether it features expressiveness based on balancing predictable and unpredictable patterns across multiple timescales. In particular, we test a songbird’s *rhythm* for evidence of such dynamic structure. The species investigated is the thrush nightingale, a songbird with a repertoire of 12-40 distinct song types per individual (a song type being a bout of continuous singing of about ~5-15s, flanked by silence) sung in immediate variety, i.e. the same song type is not usually repeated directly ^31^. Each song type contains about 4-10 different note types that have each a distinct spectral shape ^32–34^; see fig. 1A. Thrush nightingale song sounds highly “rhythmic” to human listeners ^35^,^36^, perhaps because it contains very slow as well as very fast subphrases, and many repetitions.

**Figure 1.**
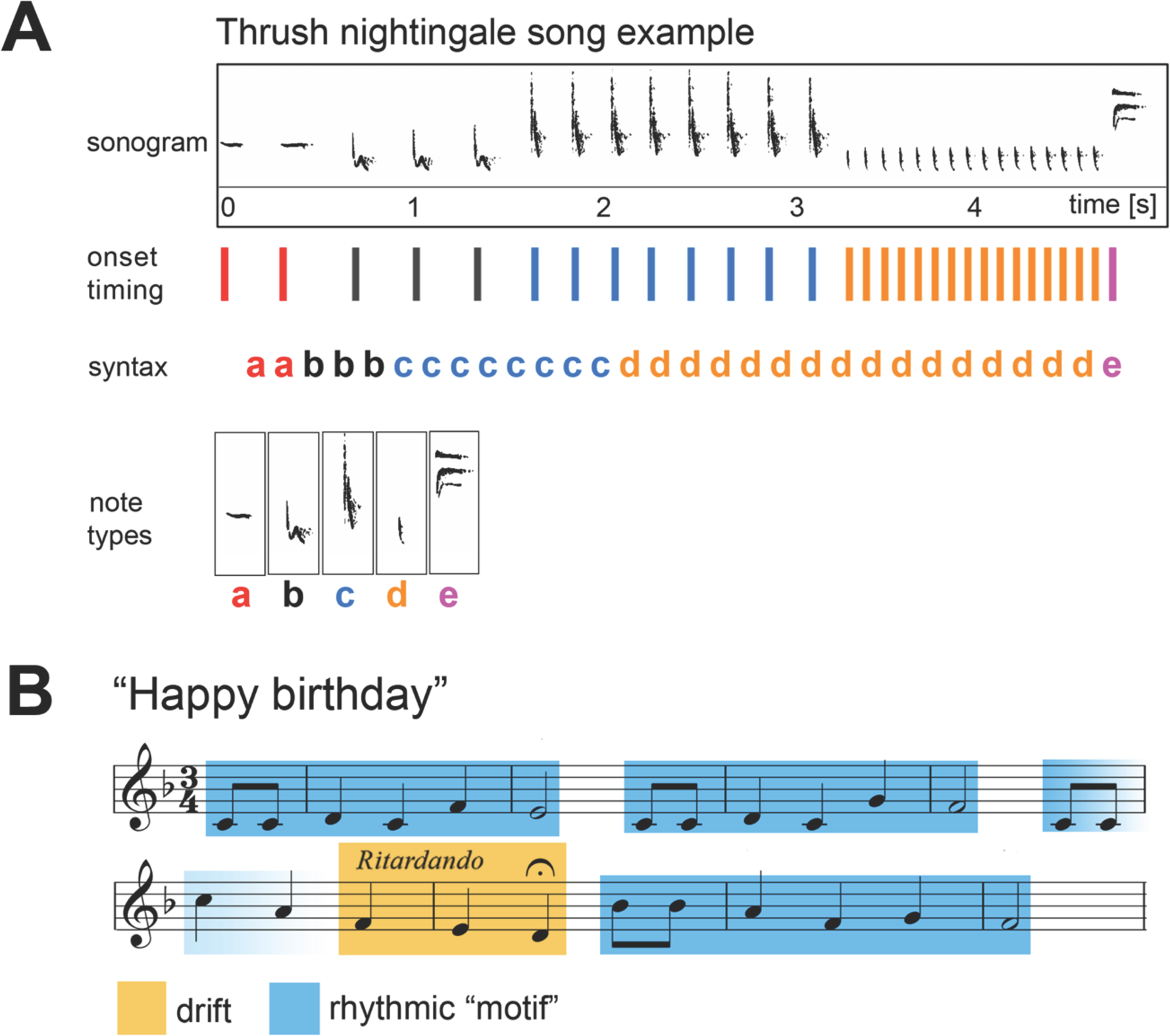
Examples illustrating rhythmic structure in music and birdsong. A,example sonogram of a thrush nightingale song consisting of five note types (a-e). The song’s rhythmic structure depends on 1) note intensity and duration (as apparent in the sonogram), 2) note syntax (i.e. sequential arrangement of notes of specific intensities and durations), and 3) timing of note onsets, which may either be under control of the bird, or an epiphenomenon of stringing together the vocal gestures required to produce the note sequence. B, score of the song “Happy birthday”, with blue shading indicating a recurring rhythmic motif (note timing in the different instances of the motif is the same, while pitch is variable), and yellow shading indicating a drift (slowing down of notes, or *ritardando* in musical terminology).

### Rhythm as note timing in birdsong

For humans, the perception of a rhythm relies on two main components: 1. the timing of single notes, and 2. the timing of accents, i.e. timing of moments of relatively higher amplitude in the note sequence ^37^.

Surprisingly little is known about songbirds’ rhythms. Rhythm, as the *timing of individual notes within the sequence* has rarely been investigated. An exception is a recent study showing that note timing in zebra finches is structured by an underlying fast, isochronous pulse that is very stable within individuals ^38^, but relative timing of notes to each other has not been systematically investigated here. This also applies to a study by Saar et al. ^39^, who developed a method to visualize how zebra finch motif length increases across development by adding more notes.

What factors could be driving note timing in the song of thrush nightingales? First, there’s the possibility that birds do not control note *timing* at all, but only the sequential arrangement of note types (syntax; see fig. 1A). Variation in note timing could merely reflect random deviations or be a trivial consequence of note choice, while the process under control of the bird would be to generate a note sequence. In that case, our human impression that thrush nightingale song sounds “rhythmic” would reflect a human perceptual bias that might not be shared by the birds, which would conceptualize their songs as sequences of elements, irrespective of internal timing. Alternatively, note timing could be under direct control of the birds. In this case, they would use sound timing and/or accentuation to generate specific rhythms. They might then – like musicians – *make use* of note timing to achieve expressiveness, driving their listeners’ expectations mixing predictable and unpredictable timing patterns.

### Expressiveness in the rhythm of music and birdsong

What does it mean that a rhythm is expressive, in the sense that its predictability/variability is fluctuating? Two strategies commonly used in music to generate such expressiveness are *recurrent patterns* and *drifts*. An example for a *recurrent pattern* in rhythm is a rhythmic motif recurring periodically within a musical sequence. The rhythm can be realized by different note types, timbres, etc., while note timing and accent within the motif are fixed. *Drifts* are successive increases or decreases in rhythmic features, like in an *accelerando* (accelerating notes) or *ritardando* (slowing down of notes). Fig. 1B shows the musical score of the song “Happy birthday”, which contains examples for both a recurrent rhythmic motif and a drift.

Both recurrences and drifts unfold across a broad range of intermediate to long timescales above transitions of individual notes, and can thus determine predictability across a (sub-)sequence of notes. For instance, during an *accelerando* phrase, the listener can predict the incoming notes to continue picking up speed, but also the acceleration to stop at some later point and lead over to a different kind of rhythm. Songbirds, like musicians, might use drifts or recurring rhythmic motifs to enable their listeners to form rhythm expectations, which can then be fulfilled, delayed, or broken.

In a thrush nightingale’s song, such expressive recurrence patterns and drifts can be generated via two different strategies: a) by *sequential arrangement* (syntax) of specific notes and pauses (fig. 2, left), or b) by adding to a given note sequence subtle *deviations* in individual note timing and/or amplitude (just like the musical operations of *accelerando, ritardando,* swinging rhythms in jazz ^40^, or *notes inégales* [unequal notes in French Baroque] would do). This way, note timing will slightly deviate from the “exact” rhythm as imposed by the sequential arrangement of note types (fig. 2, right). The two strategies have different implications for the neural mechanisms generating song: Deviation-based rhythmic structure (fig. 2, right) would require a time-shifting mechanism operating on medium to large time windows in parallel to (or upstream/downstream of) the sequence generator. Syntax based rhythmic structure (fig. 2, left) would require a neural representation of internal note features like duration and intensity, which would be used by a rhythm generating mechanism to create patterns of note timing. In either case, long memory is needed, in contrast to a situation where note timing would simply result from gesture transition without being under control by the bird. Note that a first-order model of sequence generation may still be a plausible syntax generator for sequences whose long-range structure originates in subtle deviations only (fig. 2, right), but not for sequences with long-range structure due to sequential arrangement (fig. 2, left).

**Figure 2.**
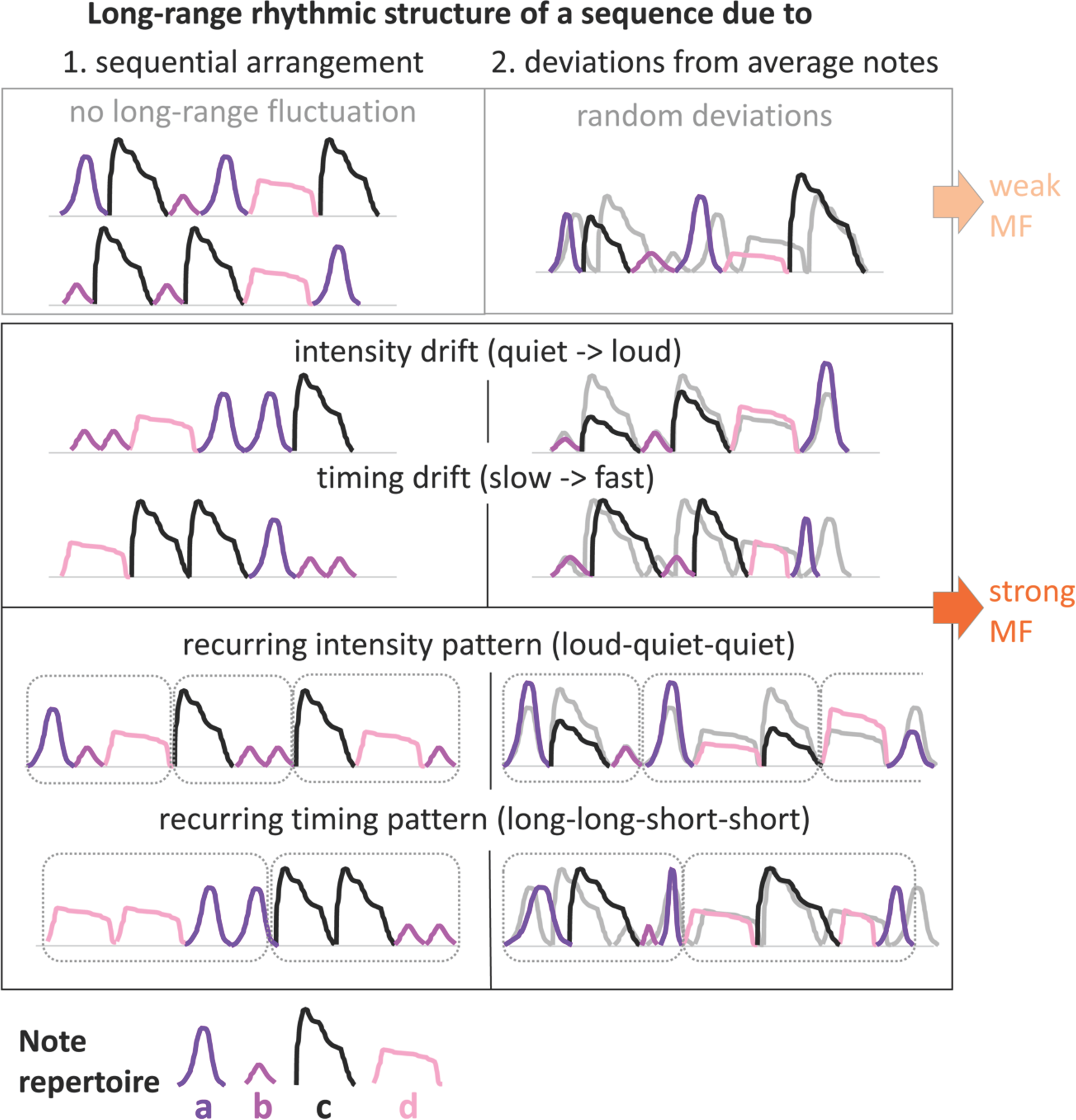
Figure 2: Impact of sequential arrangement vs. timing deviations on rhythm. Colored lines: amplitude envelopes of four notes types (a-d). Left: If sequential arrangement of notes is independent of note envelope characteristics, no systematic long-range structure emerges (top). Otherwise, arrangement may yield drifts or recurring patterns in intensity or timing (bottom). Right: Deviations of timing and amplitude from “exact” rhythm may be random (top) or generate drift or recurring patterns (bottom). Multifractality (MF) values of drifting sequences should exceed those without drift.

Music makes use of both strategies: Sequential arrangement (of notes of specific pitch and duration) has been shown to result in long-range correlations in the scores of Bach’s three-voice sinfonias ^41^, and subtle deviations from “exact” rhythm during a musical performance show long-range correlations which listeners recognize and prefer over both exact rhythm or rhythm with *random* deviations ^42,43^.

### Multifractal analysis as a test for expressiveness

Multifractal analysis is a method designed to test a signal for *fluctuations in variance across different time scales.* Hence, multifractal analysis is well suited to capture expressiveness, which we defined here as a mixture of predictable (i.e. less variable) and unpredictable (i.e. more variable) patterns across multiple timescales, and in the following we will show that multifractal analysis can be successfully applied to detect such expressiveness in birdsong.

Variability fluctuations across different timescales can be described as *changes in long-range correlations.* They way multifractal analysis uncovers such changes in long-range correlations in a signal is by calculating standard deviation over time windows of many sizes: If these changes are not random (for instance, because a pattern recurs at different times in the signal, or a drift affects multiple consecutive time windows), they have multiple fractional exponents across time (and are called “multifractal”). The larger the range of these exponents, the more does variability over time reflect interactions across multiple timescales, as opposed to merely local relationships ^44^,^45^. A birds’ rhythm that contains long-range structure, due to recurring rhythm patterns or drifts, would thus result in a wider multifractal range than note sequences lacking such features.

We first tested whether thrush nightingale rhythms exhibit such long-range correlations, by performing multifractal analysis on the songs’ amplitude envelopes (which contains information about both note timing and intensity/accentuation). Next, we tested for the role of sequential arrangement vs. timing/intensity deviations. To this end, we generated synthetic songs that lack all subtle deviations, but maintain original note order, to compare their multifractality to the original songs. To generate these “exact-rhythm” songs, we averaged for each note type the amplitude envelope across all instances, and finally re-synthesized songs from these averaged envelopes using the original *order* of notes (see Methods). These exact-rhythm songs are like a “mechanic” version of the original songs, with each note and note transition being completely stereotyped in timing and intensity. If the birds make use of timing/intensity deviations in a systematic way to increase expressiveness, the original amplitude envelopes should be more multifractal than their re-synthesized versions exact-rhythm versions.

## 2. Methods

### 2.1 Data and processing

Song data consisted in 24 thrush nightingale songs from a 2:59-min recording from the xeno-canto birdsong library (http://www.xeno-canto.org,recording #XC75409, recordist: Tomas Belka, Poland). We used “Sound Analysis for Matlab” (SAM, by Sigal Saar) to extract amplitude envelopes in 10ms windows and 1ms steps.

### 2.2 Note segmentation and identification

Several seconds separate distinct thrush nightingale songs. Within songs, we identified note boundaries by taking the difference between two Hodrick-Prescott (HP) filterings of the amplitude envelope: 1) HP filter coefficient = 50; 2) HP filter coefficient = 5*10˄7) on the 1000Hz amplitude envelope. We set between-note pauses to zero amplitude (alternate analysis showed no significant difference due to this cleaning).

### 2.3 Multifractal analysis

Multifractal analysis generalizes standard random-walk diffusion analysis that estimates how standard deviation grows as an exponent H of time ^46^. A signal like white noise, which lacks any systematic long-range structure, has a constant standard deviation as an exponent H of time (= within successively larger time windows). In contrast, a signal that contains long-range structure (like the waveform of a song with four stanzas and a chorus) contains systematic fluctuations of variability across different time windows, and its standard deviation will therefore not be uniform across time scales. To estimate the development of the standard deviation across time-scales, we proceeded as follows: First, converting series *x*(t) (in our case, a song’s amplitude envelope) of length *N* (fig. 3A) into a random walk *Y*(t) entails integration (i.e., taking cumulative sums; fig. 3B). Diffusion analysis then partitions the random-walk series into *Ns* non-overlapping windows of length *s* (10≤s≤N/4; fig. 3C-E). Linear fits *y_v_*(*t*) of random walk *Y*(t) within time windows 1≤*v*≤Ns leave mean-square residuals *F^2^* for each s:

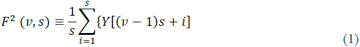

Square root of the average *F^2^* provides standard deviation *F* for each timescale s:

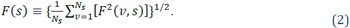

Because *H* is the exponent on time defining growth of standard deviation, diffusion analyses estimate *H* as the slope of a double-logarithmic relationship

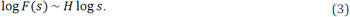

Multifractal analysis generalizes *H* with a q-order parameter that elaborates upon the squaring and square-rooting of standard deviation with qth order and qth-rooting, replacing Eq. 3 with Eq. 4 (fig. 3F) from *q_min_* to *q_max_* (in the present study, ranging from *q_min_* = 0.1 to *qmax* = 5.0):

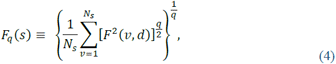

for .5<q<5.

**Figure 3.**
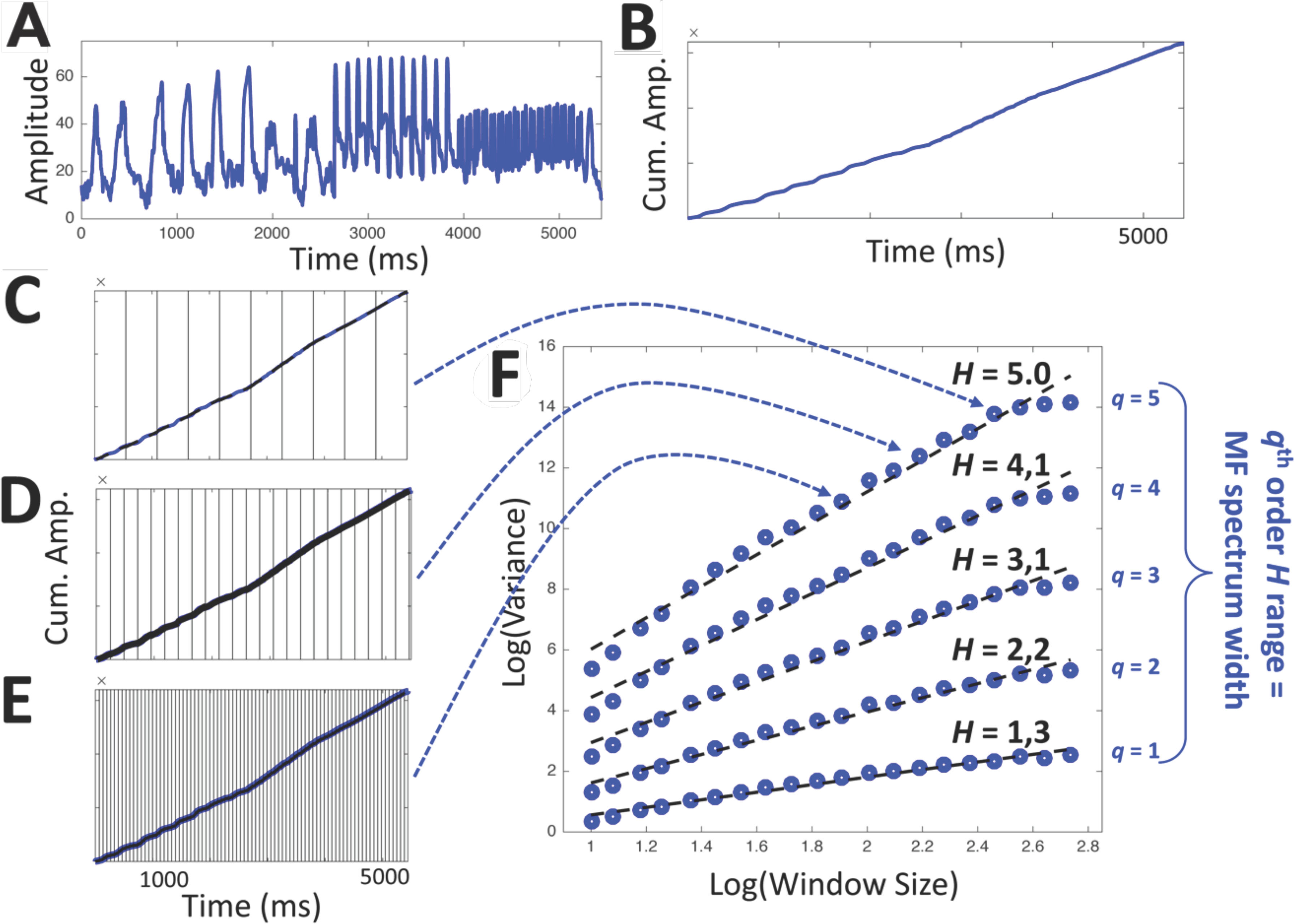
Schematic illustration of multifractal analysis. **A**, Amplitude envelope of a song and **B**, its integration. **C-E**, Three different window sizes for computing q-order deviation. **F**, Log-log plot of average *q*-order deviations vs. window size. The more long-range structure is present in a signal, the more strongly *H* differs with *q.*

### 2.5 Surrogate analysis

Spurious differences in *H* across different *q* can result from linear autocorrelation or distribution properties, without being indicative of any long-range fluctuations in variance. Hence, multifractality of the original signals are commonly compared to surrogates that preserve linear properties (i.e. distribution, linear autocorrelation) of the original signal, but lack all nonlinear structure. We compare range of qth-order *H* (multifractal spectrum width) for each original with 100 surrogate amplitude envelopes generated with the Iterative Amplitude Adjusted Fourier Transform (IAAFT) ^47^. IAAFT begins with a Fourier transform to estimate the amplitude and phase spectra. Subsequent iterations randomize the phase spectrum, applying inverse-Fourier transformation to combine it with the original amplitude spectrum, and then replace rank-ordered values of this new series with those of the original series.

Evidence for systematic variance fluctuations/long-range structure across hierarchically nested timescales is a multifractal spectrum width for the signal in question that lies beyond the 95% confidence interval its IAAFT surrogates ^44^.

### 2.6 Testing synthetic “exact-rhythm” songs

We classified note types by visual and auditory inspection using SoundAnalysisPro and GoldWave v6.18. For each type, we created an envelope profile by averaging note duration and intensity course across all instances. Pause types (as identified by the two adjacent note types) were averaged for duration (pause amplitude being set to zero). Using these averaged note and pause profiles, we generated synthetic songs using the original sequential arrangement. Multifractal spectrum widths of these “exact rhythm” songs were then compared to their IAAFTs like the original songs.

## 3 Results

To identify any systematic long-range variability fluctuations in the thrush nightingale rhythms, we first calculated multifractality of the original songs’ amplitude envelopes, and compared this value to the multifractality value of their IAAFT surrogates (control time series with the same distribution and linear autocorrelation properties but lacking any long-range correlations; see Methods). All 24 songs’ amplitude envelopes exhibited multifractality, i.e. non-zero ranges in the qth-order *H* (fig. 4A). Further, multifractal spectrum widths for original amplitude envelopes (black circles) are above the 95%-confidence intervals of their corresponding IAAFT surrogates. This indicates that song amplitude envelopes exhibit non-local changes in variability, systematically going through more and less variable phases across different timescales.

We next compared the original rhythms to the “exact” rhythms of songs that we stripped off any subtle timing and amplitude deviations, to test whether birds add expressiveness to their vocal sequences by adding systematic timing/intensity fluctuations. The exact-rhythm songs turned out still more multifractal than their IAAFT controls, but less so than the original songs: A t-test revealed that the originals are significantly more multifractal (i.e. differed more strongly from their IAAFT) compared to their matched exact-rhythm songs (fig. 4B; t(23) = 8.41, *p* < .001). Thus, a significant part of the original rhythms’ multifractality originates in non-random deviations from average note/pause profiles. Another part of the multifractality – the part still present in the exact-rhythm songs – is due to the sequential arrangement of notes. Drifts and motif recurrences that contributed to the rhythms’ multifractality can be apparent in the sonograms (fig. 4C).

**Figure 4:**
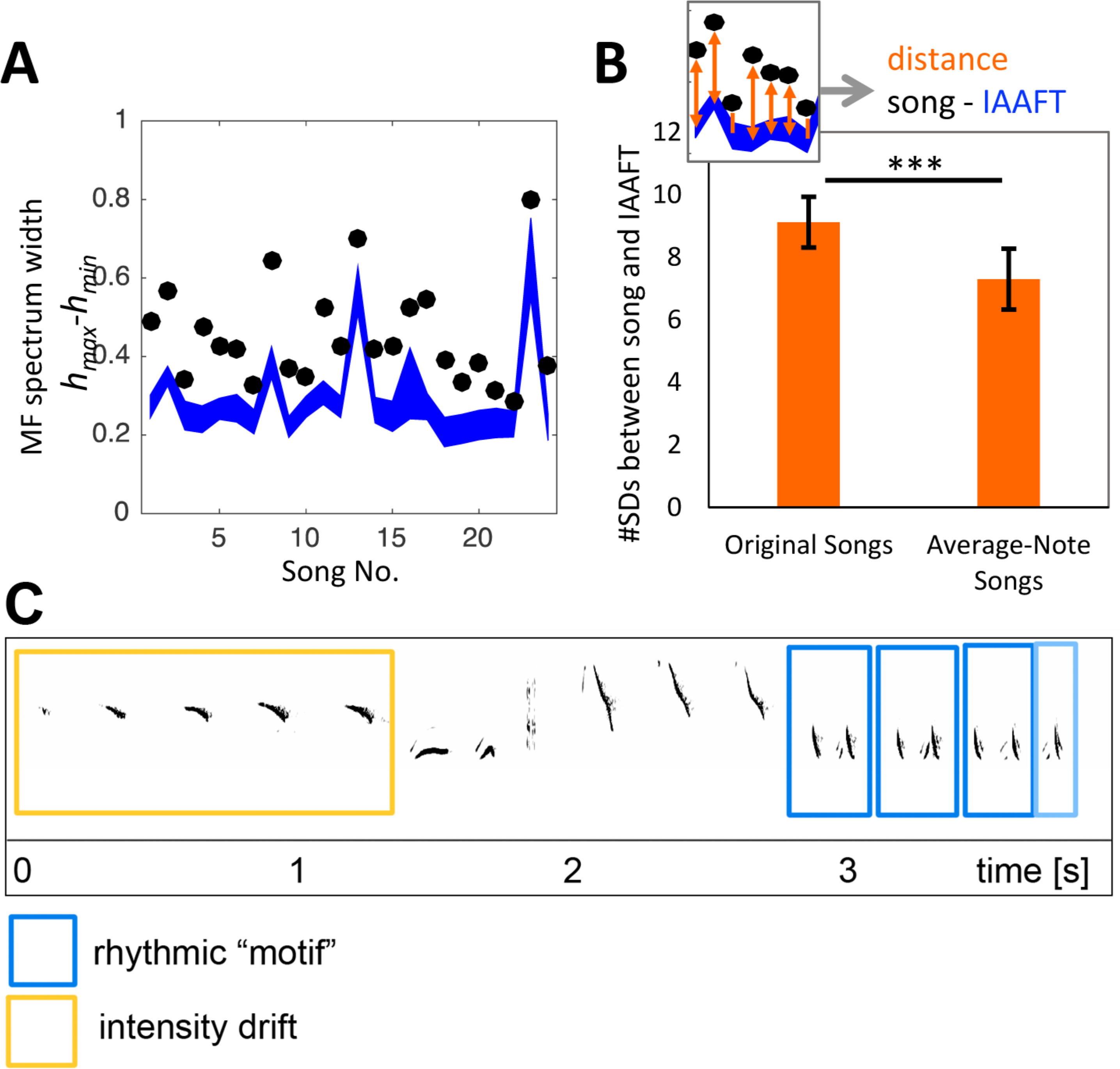
Thrush nightingale rhythms are multifractal, due to both sequential arrangement and timing deviations. **A**, Multifractal spectra of the 24 original songs (black circles) are wider than expected from their IAAFT surrogates (blue). Blue area = 95% confidence interval of the surrogates. **B**, Multifractality in original rhythms is significantly greater compared to rhythm of “exact rhythm” songs. Bars represent *difference* between amplitude envelopes and their IAAFTs (inset), measured in *standard deviations from the mean* of the IAAFTs' multifractality values *(hmax-h_m_m).* Averaging note amplitude envelopes significantly reduced multifractality (p < .001). **C**, Thrush nightingale song containing examples of an intensity drift (yellow) and a rhythmic motif (blue).

## 4 Discussion

Using multifractal analysis, we show that the rhythms sung by a thrush nightingale contain long-range correlations across multiple timescales, and that part of this structure is due to subtle timing and intensity deviations from average note/pause profiles: Eliminating those subtle deviations resulted in significantly reduced multifractality, indicating that these deviations are not random. Instead, they contribute to systematic long-range correlations in the songs’ rhythm – for instance, by adding drifts or recurrent rhythm patterns to the note sequence. However, songs stripped off all these subtle deviations were still significantly more multifractal than their IAAFT controls. This indicates that another part of the rhythms’ multifractality is due to sequential arrangement of the particular note types: By combining notes with specific internal structures into sequences, the birds also generate long-range rhythm patterns, which may materialize as drifts or recurrences.

Our results are the first to show multifractality in birdsong, and show that multifractal analysis can be successfully used to determine the extent of structuredness in animal vocalizations.

The multifractality of the thrush nightingale rhythm strongly suggests that note timing is under the control of the birds, instead of merely being an epiphenomenon of syntax, i.e. only constrained by the peripheral dynamics of the vocal apparatus producing a sequence of particular gestures.

Our findings suggest that thrush nightingales produce rhythms that are more complex than what a simple Markovian model of sequence generation could specify. The rhythms are not exhaustively conceptualized by a finite number of independent processes, such as drawing from a distribution of notes, and stringing them up into longer sequences by an independent combination process using local rules ^48^. Instead, the generating mechanism to produce such sequences requires a) memorizing song features trailing back more than just one or a few notes, b) taking into account the discrete identity *and* internal features of the note types sung (i.e. their duration, intensity, amplitude course), and c) being able to manipulate note onset times and intensities with a time/intensity-shifting mechanism that operates on the time-scales of (sub-)phrases and is independent of operations regulating the sequential arrangement of notes. Such a mechanism might be analogous to the processing streams in the human brain assumed to process prosody (sentence melody): These are separate from and working in parallel to the processing streams that underlie core linguistic abilities such as phonology, syntax, and semantics ^49^.

In addition to such parallels to language processing, our findings underline similarities between birdsong and music structure. Long-range correlations similar to the ones we found in the thrush nightingale rhythms have also been described for musical rhythm: “Exact” rhythm (as noted in the score) has been shown to be multifractal in classical pieces ^50^, analogous to the multifractality we find in “exact rhythm” songs. Moreover, human listeners prefer multifractal timing deviations from the beat ^42^,^43^, similar to the significant contribution of timing/intensity deviations to multifractality in our data. Multifractality in note arrangement and timing deviations might reflect expressiveness intentionally added by birds and humans for attractiveness. Further experimental research is needed to explore whether avian rhythms with higher multifractality are more effective in engaging their listeners’ attention.

In sum, our results support the hypothesis that songbirds and musicians use similar techniques of combining expected and unexpected patterns to attract their listeners’ attention. Two mechanisms to do so are 1) arranging notes and 2) adding expressive timing to a song in a way that long-range rhythm structure emerges, for example as recurrent timing/intensity patterns or drifts.

## Data accessibility

Data is accessible from the xeno-canto birdsong library (for details, see Methods 2.1)

## Ethics

n/a.

## Author contributions

Conceptualization, Methodology/Analysis T.R., S.W; Writing of manuscript, Preparation, Review & Editing, Final Approval: T.R., S.W., and D.K.-S. Figure preparation, Review & Editing, Final Approval: T.R., S.W., and D.K.-S.

## References

1. Morton, E. S. Ecological Sources of Selection on Avian Sounds. Am. Nat. 109, 17–34 (1975).

2. Marten, K. & Marler, P. Sound transmission and its significance for animal vocalization - I. Temperate habitats. Behav. Ecol. Sociobiol 2, 271–290 (1977).

3. Marten, K., Quine, D. & Marler, P. Sound transmission and its significance for animal vocalization - II. Tropical forest habitats. Behav. Ecol. Sociobiol. 2, 291–302 (1977).

4. Wiley, R. & Richards, D. Physical constraints on acoustic communication in the atmosphere: Implications for the evolution of …. Behav. Ecol. Sociobiol. (1978).

5. Cosens, S. E. & Falls, J. B. A comparison of sound propagation and song frequency in temperate marsh and grassland habitats. Behav. Ecol. Sociobiol. 15, 161–170 (1984).

6. Sorjonen, J. Factors affecting the structure of song and the singing behavior of some northern European passerine birds. Behaviour 98, 286–304 (1986).

7. Sorjonen, J. Transmission of the Two Most Characteristic Phrases of the Song of the Thrush Nightingale Luscinia luscinia in Different Environmental Conditions. Ornis Scand. (Scandinavian J. Ornithol. 14, 278–288 (1983).

8. Brumm, H. The impact of environmental noise on song amplitude in a territorial bird. J. Anim. Ecol. 73, 434–440 (2004).

9. Boncoraglio, G. & Saino, N. Habitat structure and the evolution of bird song: A meta-analysis of the evidence for the acoustic adaptation hypothesis. Funct. Ecol. 21, 134–142 (2007).

10. Naguib, M. Reverberation of rapid and slow trills: Implications for signal adaptations to long-range communication. J. Acoust. Soc. Am. 113, 1749–1756 (2003).

11. Brumm, H. & Naguib, M. Chapter 1 Environmental Acoustics and the Evolution of Bird Song. Advances in the Study of Behavior 40, 1–33 (2009).

12. Brumm, H. Signalling through acoustic windows: Nightingales avoid interspecific competition by short-term adjustment of song timing. J. Comp. Physiol. A Neuroethol. Sensory, Neural, Behav. Physiol. 192, 1279–1285 (2006).

13. Griessmann, B. & Naguib, M. Song sharing in neighboring and non-neighboring thrush nightingales (Luscinia luscinia) and its implications for communication. Ethology 108, 377–387 (2002).

14. Naguib, M. Effects of song overlapping and alternating on nocturnally singing nightingales. Anim. Behav. 58, 1061–1067 (1999).

15. Bhattacharya, H., Cirillo, J. & Todt, D. Universal features in the singing of birds uncovered by comparative research. Our Nat. 6, 1–14 (2008).

16. Hasselquist, D. & Bensch, S. Daily energy expenditure of singing great reed warblers acrocephalus arundinaceus. J. Avian Biol. 39, 384–388 (2008).

17. Eberhardt, L. S. Oxygen consumption during singing by male carolina wrens (Thryothorus ludovicianus). Auk 111, 124–130 (1994).

18. Gil, D. & Gahr, M. The honesty of bird song: Multiple constraints for multiple traits. Trends in Ecology and Evolution 17, 133–141 (2002).

19. Thomas, R. J. The costs of singing in nightingales. Anim. Behav. 63, 959–966 (2002).

20. Podos, J. A Performance Constraint on the Evolution of Trilled Vocalizations in a Songbird Family (Passeriformes: Emberizidae). Evolution (N. Y). 51, 537 (1997).

21. ten Cate, C. & Okanoya, K. Revisiting the syntactic abilities of non-human animals: natural vocalizations and artificial grammar learning. Philos. Trans. R. Soc. London B Biol. Sci. 367, (2012).

22. Berwick, R. C., Okanoya, K., Beckers, G. J. L. & Bolhuis, J. J. Songs to syntax: the linguistics of birdsong. Trends Cogn. Sci. 15, 113–21 (2011).

23. Kershenbaum, A. et al. Animal vocal sequences: not the Markov chains we thought they were. Proc. R. Soc. B Biol. Sci. 281, 20141370–20141370 (2014).

24. Rothenberg, D., Roeske, T. C., Voss, H. U., Naguib, M. & Tchernichovski, O. Investigation of musicality in birdsong. Hear. Res. 308, 71–83 (2014).

25. Huron, D. Sweet anticipation: Music and the psychology of expectation. (2006).

26. Meyer, L. Emotion and meaning in music. 1956. …. Affect. Exp. apprehension Music. … (1956).

27. Levitin, D. This is your brain on music: The science of a human obsession. (2006).

28. Cross, I. & Narmour, E. The Analysis and Cognition of Melodic Complexity. (1995).

29. de Mántaras, R. L. & Arcos, J. L. AI and music: From composition to expressive performance. AI Mag. 23, 43–58 (2002).

30. Broomhead, P. Shaping Expressive Performance: A Problem-Solving Approach. Music Educ. J. 91, 63 (2005).

31. Naguib, M. & Todt, D. Recognition of neighbors’ song in a species with large and complex song repertoires: the Thrush Nightingale. J. Avian Biol. 29, 155–160 (1998).

32. Naguib, M. & Kolb, H. COMPARISON OF THE SONG STRUCTURE AND SONG SUCCESSION IN THE THRUSH NIGHTINGALE (LUSCINIA-LUSCINIA) AND THE BLUE THROAT (LUSCINIA-SVECICA). J. FUR Ornithol. 133, 133–145 (1992).

33. Lille, R. Art-und Mischgesang von Nachtigall und Sprosser (Luscinia megarhynchos, L. luscinia). J. für Ornithol. 129, 133–159 (1988).

34. Sorjonen, J. Seasonal and diel patterns in the song of the thrush nightingale Luscinia luscinia in SE Finland. Ornis Fenn. (1977).

35. Rothenberg, D. Why birds sing: A journey into the mystery of bird song. (2006).

36. Sotavalta, O. Song patterns of two sprosser nightingales. Ann. Finnish Zool. Soc. (1956).

37. Patel, A. D.. & Lyon, B. Music, Language, and the Brain by Aniruddh D. Patel. Psychomusicology 20, 182–187 (2008).

38. Norton, P. & Scharff, C. ‘Bird Song Metronomics’: Isochronous Organization of Zebra Finch Song Rhythm. Front. Neurosci. 10, 309 (2016).

39. Saar, S. & Mitra, P. P. A technique for characterizing the development of rhythms in bird song. PLoS One 3, (2008).

40. Friberg, A. & Sundström, A. Swing Ratios and Ensemble Timing in Jazz Performance: Evidence for a Common Rhythmic Pattern. Music Percept. 19, 333–349 (2002).

41. Telesca, L. & Lovallo, M. Revealing competitive behaviours in music by means of the multifractal detrended fluctuation analysis: application to Bach’s Sinfonias. Proc. R. Soc. London A Math. Phys. Eng. Sci. 467, (2011).

42. Räsänen, E. et al. Fluctuations of Hi-Hat Timing and Dynamics in a Virtuoso Drum Track of a Popular Music Recording. PLoS One 10, e0127902 (2015).

43. Hennig, H. et al. The Nature and Perception of Fluctuations in Human Musical Rhythms. PLoS One 6, e26457 (2011).

44. Ihlen, E. Introduction to multifractal detrended fluctuation analysis in Matlab. Fractal Anal (2012).

45. Kelty-Stephen, D., Palatinus, K. & Saltzman, E. A tutorial on multifractality, cascades, and interactivity for empirical time series in ecological science. Ecological (2013).

46. Kantelhardt, J. & Zschiegner, S. Multifractal detrended fluctuation analysis of nonstationary time series. Phys. A Stat. (2002).

47. Schreiber, T. & Schmitz, A. Improved surrogate data for nonlinearity tests. Phys. Rev. Lett. (1996).

48. Kelty-Stephen, D. G. & Wallot, S. Multifractality versus (mono) fractality in the narrative about nonlinear interactions across time scales: Disentangling the belief in nonlinearity from the diagnosis of nonlinearity. Ecol. Psychol. (2017).

49. Sammler, D., Grosbras, M. H., Anwander, A., Bestelmeyer, P. E. G. & Belin, P. Dorsal and ventral pathways for prosody. Curr. Biol. 25, 3079–3085 (2015).

50. Su, Z.-Y. & Wu, T. Multifractal analyses of music sequences. Phys. D Nonlinear Phenom. 221, 188–194 (2006).

